# VRS5 (HvTB1) binds to the promoter of tillering and floral homeotic genes to regulate their expression

**DOI:** 10.1101/2024.08.22.609225

**Authors:** T. Winkelmolen, P. Colleoni, M. Moscou, P. Hoseinzadeh, K. Oldach, R.C. Schmidt, R. Immink, G.W. van Esse

## Abstract

Variation in shoot architecture, or tillering, is an important adaptive trait targeted during the domestication of crops. A well-known regulatory factor in shoot architecture is TEOSINTE BRANCHED 1 (TB1). TB1 and its orthologs have a conserved function in integrating environmental signals to regulate axillary branching, or tillering in cereals. The barley ortholog of TB1, *VULGARE ROW-TYPE SIX 5* (VRS5) does not only regulate tillering, but is also involved in regulating row-type by inhibiting lateral spikelet development. These discoveries predominantly come from genetic studies, but how VRS5 regulates these processes on a molecular level remains largely unknown. By combining transcriptome analysis between *vrs5* and wild type at different developmental stages and DAP-sequencing to locate the genome-wide DNA binding sites of VRS5, we identified *bona fide* targets of VRS5. We found that VRS5 targets abscisic acid related genes, potentially to inhibit tillering in a conserved way. While later in inflorescence development, row-type gene *VRS1* and several known floral development genes, like MIKCc type MADS-box genes, are targeted. This study identifies several significant genes for mutational analysis, representing a selection of *bona fide* targets that will contribute to a deeper understanding of the VRS5 network and its role in shaping barley development.

## Introduction

Variation in shoot architecture is one of the most important adaptive traits that have been targeted during crop domestication (Doebley et al., 2006). In cereals like barley and wheat, the aboveground architecture is determined by the number of branches, called tillers, and inflorescence (spike) architecture (Dixon et al., 2022). Tillers are formed during vegetative growth of the shoot apical meristem, which produces successive leaves with (vegetative) axillary meristems (AM) in the leaf axil. The regulation of tillering usually happens after AM initiation, where the developing AM can enter a stage of dormancy as a bud (Hussien et al., 2014; Riaz et al., 2023; Shang et al., 2021). The trigger to develop a bud into a tiller, or to remain dormant, depends on both internal and external cues (Dixon et al., 2022; Shaaf et al., 2019; Wang et al., 2018).

One of the key players that is involved in the control of bud outgrowth is TEOSINTE BRANCHED 1 (TB1). TB1 orthologs have been targeted during domestication for their impact on plant architecture and development (Clark et al., 2004; Martín-Trillo et al., 2011; Wang et al., 1999). In maize for example, increased expression of *ZmTB1* contributes to a complete suppression of the lateral tillers and increased crop yield (Doebley et al., 1995; Studer et al., 2011). TB1 is a member of the TEOSINTE BRANCHED 1/CINCINNATA/PROLIFERATING CELL FACTOR (TCP) family. TCP family members regulate gene expression trough DNA binding and act as transcriptional regulators. Recent studies in maize indicate that at the molecular level TB1 directly regulates genes involved in hormone signaling, including abscisic acid (ABA), gibberellic acid (GA), and jasmonate (JA), and sugar signaling (Dong et al., 2019).

Besides its function in the control of tiller outgrowth, TB1 plays an additional role in orchestrating spike architecture in the *Triticeae*. In wheat, high levels of *TaTB1-D1* result in reduced expression of meristem identity genes, thereby promoting paired spikelet formation (Dixon et al., 2018). In barley, the *TB1* ortholog, *VULGARE ROW-TYPE SIX 5* (*VRS5*), which is also known as *INTERMEDIUM-C* (*INT-C*), impacts not only tiller development, but also represses lateral spikelet development. When compared to wild-type lines, *vrs5 (int-c)* mutants exhibit an increased tiller number at early developmental stages and developed lateral florets of the triple spikelet meristem, leading to a six-rowed spike phenotype in contrast to a two-rowed spike in wild-type barley (Liller et al., 2015; Ramsay et al., 2011; Zwirek et al., 2019). In two-rowed cultivars, the formation of lemma and carpel primordia stagnates in the lateral florets, making them infertile, which means the lateral florets do not form a grain. In contrast, carpel primordia are formed and fully developing in *vrs5* mutants, making them fertile, which means the lateral florets do develop into grains (Zwirek et al., 2019). Taken together, functional VRS5 in 2-rowed cultivars, prevents the formation of carpel primordia in the lateral spikelets. In line with the mode of action of VRS5 in tiller and lateral spikelet development, *VRS5* is expressed in the tiller buds and the lateral spikelet primordia of developing inflorescences (Thiel et al., 2021; Wang et al., 2022).

Despite the key role of VRS5 in orchestrating tillering and lateral spikelet development in barley, the molecular mode of action remains poorly understood. To gain a better understanding of the regulatory network by which VRS5 controls both tiller and inflorescence development, we have profiled transcriptional changes associated with altered development in the *vrs5* mutant. This comprehensive analysis enabled us to define the *vrs5*-related transcriptional changes over development. In combination with a genome-wide transcription factor (TF) binding site mapping using DNA affinity purification (DAP-seq), we identified *bona fide* VRS5 targets. Amongst these *bona fide* targets are floral homeotic and meristem identity genes. In addition, our transcriptional profiling combined with DAP-seq using VRS5 as bait, reveals an evolutionary conserved and divergent role of VRS5 in the regulation of tiller and grain development. Taken together, this study shows a comprehensive view of the core molecular network downstream of VRS5 and provides novel candidate genes that can be exploited in the regulation of tillering and inflorescence architecture in cereals.

## Materials and Methods

### Phenotypic analysis

Seeds of Bowman and *int-c.5* (GSHO 2003) were stratified on wet tissue paper in the dark at 4°C for three days, before sowing them in 1.3L pots. Plants were grown in the greenhouse under 16h light 21°C, 8h dark 18°C conditions. Tiller number was counted per plant at 7, 10, and 14 days after emergence (DAE).

### RNA isolation and RNA-Sequencing

Seeds of cv. Bowman and *int-c.5* were sown in 84-well trays. To improve equal germination, a stratification treatment was provided by exposing the seeds at 4°C in the dark for three days. After stratification, plants were transferred to soil in a growth chamber under long day (LD) conditions (16h light/ 8h dark) at 20°C. Developmental stages of the meristems were scored using the Waddington scale (WADDINGTON et al., 1983) by dissecting three plants of each genotype on the day of sampling. Shoot apical meristems were sampled over time at four different stages: W0, vegetative apex (VA); W1, transition apex (TA); W2.25, triple mound (TM); and W3-3.5, lemma and stamen primordia (LP/SP). Samples were taken in the afternoon (6 hours before the end of the LD period). Leaves surrounding the apex were removed manually, before cutting the apex. Apexes were immediately frozen in liquid nitrogen and stored at −80°C before RNA isolation. Frozen tissue was disrupted using the Tissuelyser LT (Qiagen). RNA extraction was performed on disrupted tissues using columns from the GeneJET plasmid miniprep kit (Thermo Fischer Scientific), as described previously (Yaffe et al., 2012). Extracted RNA was treated with DNase I (Qiagen) and RiboLock (Fermentas).

Extracted RNA samples that were used for RNA-Seq (Novogene Europe, Cambridge Science Park, United Kingdom), where mRNA was purified from total RNA using poly-T oligo-attached magnetic beads. After fragmentation, the first strand cDNA was synthesized using random hexamer primers, followed by the second strand cDNA synthesis. Samples were sequenced using the Illumina platform with a sequencing depth of 20 million reads per sample using 150 bp paired-end reads.

### Differential gene expression analysis

Raw paired RNA-seq reads were cleaned using Trimmomatic 0.39 (Bolger et al., 2014) with parameters: ILLUMINACLIP:adapters.fa:2:30:10 LEADING:3 TRAILING:3 SLIDINGWINDOW:4:15 MINLEN:36. The file adapters.fa contains nucleotide sequences of Illumina adapters used for library construction. From the trimmed reads, transcripts were quantified with the MorexV3 transcriptome as reference (Mascher et al., 2021) using salmon (Patro et al., 2017) and the softclip parameter. Differentially expressed genes were identified using the R package DESeq2 (Love et al., 2014) per developmental stage (VA, TA, TM, and LP/SP), and over development by adding developmental stage to the model (GLM). Identified differentially expressed genes were considered significant when the false discovery rate (FDR) < 0.05 and the absolute log2 fold change (LFC) was more than 0.58. Data from the later developmental stages, namely the stamen primordia stage (SP), and the awn primordia stage (AP), when the carpels are formed and the first phenotypical differences between Bowman and *int-c.5* in spike development emerge, are extracted from Gene Expression Omnibus (GEO) expression data set GSE102191 (van Esse et al., 2017). The data from these stages were re-analyzed against the Morex V3 genome in the same way as the newly generated RNA-Seq samples from the VA, TA, TM and LP/SP developmental stages.

### Gene Ontology (GO) term annotation, enrichment and cluster analysis

Using Pannzer2 (Törönen et al., 2018), the coding sequences of MorexV3 reference genome (Mascher et al., 2021) were annotated and assigned GO-terms (accessed 21-06-2023). Genes that were significant differentially expressed in at least one developmental stage and/or independent of developmental stage were clustered based on their expression pattern per genotype using the degPatterns function from the R package DEGreport. In total, 2,869 transcripts show differential expression with |Log2FoldChange| > 0.58 and FDR < 0.05 in at least one of the developmental stages or independent of developmental time in the GLM. Clusters were grouped manually based on their global trend over time in six cluster groups. GO-term enrichment of the differentially expressed genes was done for each cluster group using the R package topGO using the “classic” algorithm as parameter followed by the Fisher’s Exact Test, taking into account only those GO-terms that have at least 10 annotated genes in the designated term.

### DNA affinity purification and sequencing (DAP-seq)

DNA libraries for DAP-Seq were made as previously described (Bartlett et al., 2017). For each replicate, genomic DNA was extracted from 100 mg developing inflorescences of barley cv. Morex, four weeks after sowing in 84-well trays (Waddington stage ∼W4) using the DNeasy Plant Mini kit (Qiagen). The DNA was fragmented with ten sonication steps of ten seconds using the Sonicator 150 (MSE) and allowing 45 seconds of rest on ice between each step. The fragmented DNA was end-repaired using the End-It kit (Lucigen) and A-tailed using Klenow fragment (3’-5’ exo-; NEB). The truncated Illumina Y-adapter was ligated to the DNA using T4 DNA ligase (Promega).

The CDS of VRS5 (cv. Bowman) without the stop codon was cloned, via Golden Gate cloning, together with a C-terminal 3xFLAGtag (MoClo toolbox; pICSL50007) (Engler et al., 2014) behind the SP6 promoter into the pSPUTK expression vector (Stratagene) that was made Golden Gate compatible as described (Kerstens et al., 2024). Similarly, eGFP was cloned from pICH41531. The FLAG-tagged VRS5 and GFP were expressed in-vitro using 2 µg plasmid and TnT® SP6 High-Yield Wheat Germ Protein Expression System (Promega). Expressed FLAG-tagged proteins were bound with prepared DNA libraries for 2 hours at RT followed by immobilization by anti-FLAG magnetic beads (SIGMA). The mix was washed, and eventually DNA was eluted by competition with 3xFLAG peptide (APExBIO). Eluted DNA was amplified by PCR with primers completing the Illumina sequencing adapter of which one primer was indexed (**Table S1**). Amplified products were run on a 1.5% agarose gel, and DNA between 200 and 600 bp was cut out and purified using GeneJET Gel Extraction Kit (Thermo Scientific). Six replicates (**Figure S1**) were sequenced using the HiSeq X-ten platform with a sequencing depth of 50 million reads per sample using 150 bp paired-end reads (BGI Europe).

### DAP-seq analysis

Raw paired DAP-seq reads were cleaned using Trimmomatic 0.39 (Bolger et al., 2014) using parameters ILLUMINACLIP:adapters.fa:2:30:10 LEADING:3 TRAILING:3 SLIDINGWINDOW:4:15 MINLEN:36 (**Supplemental Table 1**). Paired reads were mapped to the Barley MorexV3 reference genome (Mascher et al., 2021) using BWA-MEM v0.7.17 (Li, 2013, preprint; Li and Durbin, 2010). Peak calling was done using MACS2 v2.2.7.1 (Zhang et al., 2008), using the callpeak function, where the VRS5 samples were compared to the GFP control samples. The genome size parameter was set to 5.4e9, corresponding to the barley genome size. The “KEEP-DUP” parameter was set to “auto”, meaning that the MACS2 algorithm calculates the number of reads that are allowed to be identical, leading to a maximum of two identical reads. Genomic regions that had a high number of reads in the control samples (GFP-FLAG) were identified using the “greenscreen” method (Klasfeld et al., 2022). Called peaks overlapping with a “greenscreen” region were filtered out using bedtools intersect function (Quinlan and Hall, 2010). Peaks with a q < 10^-10^ were kept and considered true peaks. The DNA sequences of the true peaks were used for de novo motif discovery using STREME from MEME-suite v5.4.1 (Bailey, 2021; Bailey et al., 2015). The bdg files from MACS2 output were converted to tdf files for visualization using the Integrative Genomics Viewer v.2.11.9 (Robinson et al., 2011).

### Overlap with ZmTB1 targets

To check if previously identified ZmTB1 targets (Dong et al., 2019) are also targeted by VRS5, we used Orthofinder (Emms and Kelly, 2019) to infer the barley orthologs of the ZmTB1 targets. As input the following proteomes are downloaded from Phytozome: *Arabidopsis thaliana* TAIR10 (Lamesch et al., 2012), *Brachypodium distachyon* v3 (Vogel et al., 2010), *Hordeum vulgare* (barley) MorexV3 (Mascher et al., 2021), *Lolium perenne* MPB_Lper_Kyuss_1697 (Frei et al., 2021), *Triticum aestivum* (wheat) IWGSC (ENA: GCA_900519105.1), *Oryza sativa* (rice) IRGSP 1.0 (Ouyang et al., 2007), and *Zea mays* (maize) AGPv3.31 (NCBI: GCF_000005005.1), which was used as reference by Dong et al., 2019.

#### CRISPR-Cas9 mutagenesis

For CRISPR-Cas9 targeted mutagenesis of *VRS1*, three guides were cloned using a *TaU6* as promoter and an optimized scaffold design for improving CRISPR-Cas9 efficiency (Dang et al., 2015). The guides include guide 1: 5’-ACGTGGACACGACTTTCTTC-3’, guide 2: 5’-GAGGAGGGGACCCCAAGAAG-3’, and guide 3: 5’-GGAGGTGCGGCGCCTCAGGT-3’. Modules were cloned using the Golden Gate vector system (Engler et al., 2014). The sequence-verified construct was introduced in hypervirulent *Agrobacterium tumefaciens* strain AGL1 pSOUP. Transformation of barley plants was done as previously described (Hensel et al., 2009). T_0_ plants were genotyped for CRISPR cuts using the Phire Plant Direct PCR Kit (Thermo Fisher Scientific) using primers described in **Table S1**. Amplified fragments were sequenced directly.

### Data availability

The data generated in this publication have been deposited in NCBI’s Gene Expression Omnibus (Barrett et al., 2013; Edgar et al., 2002) and are accessible through GEO Series accession number GSE272367 and GSE272368.

## Results

### VRS5 regulates both tillering and row-type

Previous mutant analysis revealed a role for *VRS5* in tillering and in determination of the row-type in the spikelet (Liller et al., 2015; Ramsay et al., 2011; Wang et al., 2022). Differences in tillering between *vrs5* and Bowman are observed already early in development, when the meristem is still vegetative (**Figure 1A+B, Figure S2A**). To identify the genes regulated by VRS5, we generated transcriptomes of WT (Bowman) and *vrs5 (int-c.5)* mutant shoot apexes at four different developmental stages. By doing a transcriptome analysis over four developmental stages, we aim to distinguish between the role of VRS5 on tillering and on inflorescence development. We took the vegetative apex (VA) and transition apex (TA) as two early developmental stages (**Figure 1B**). The entire crown was used to ensure that emerging tillers containing tiller buds were sampled along with the meristem. To capture the genes differentially regulated early in floral organ development we isolated RNA for transcriptome profiling from the triple mound (TM), and lemma and stamen primordia stage (LP/SP), respectively. At these stages, the central and lateral floret primordia are initiated, and the first floral organ primordia are formed (**Figure 1B**).

**Figure 1:**
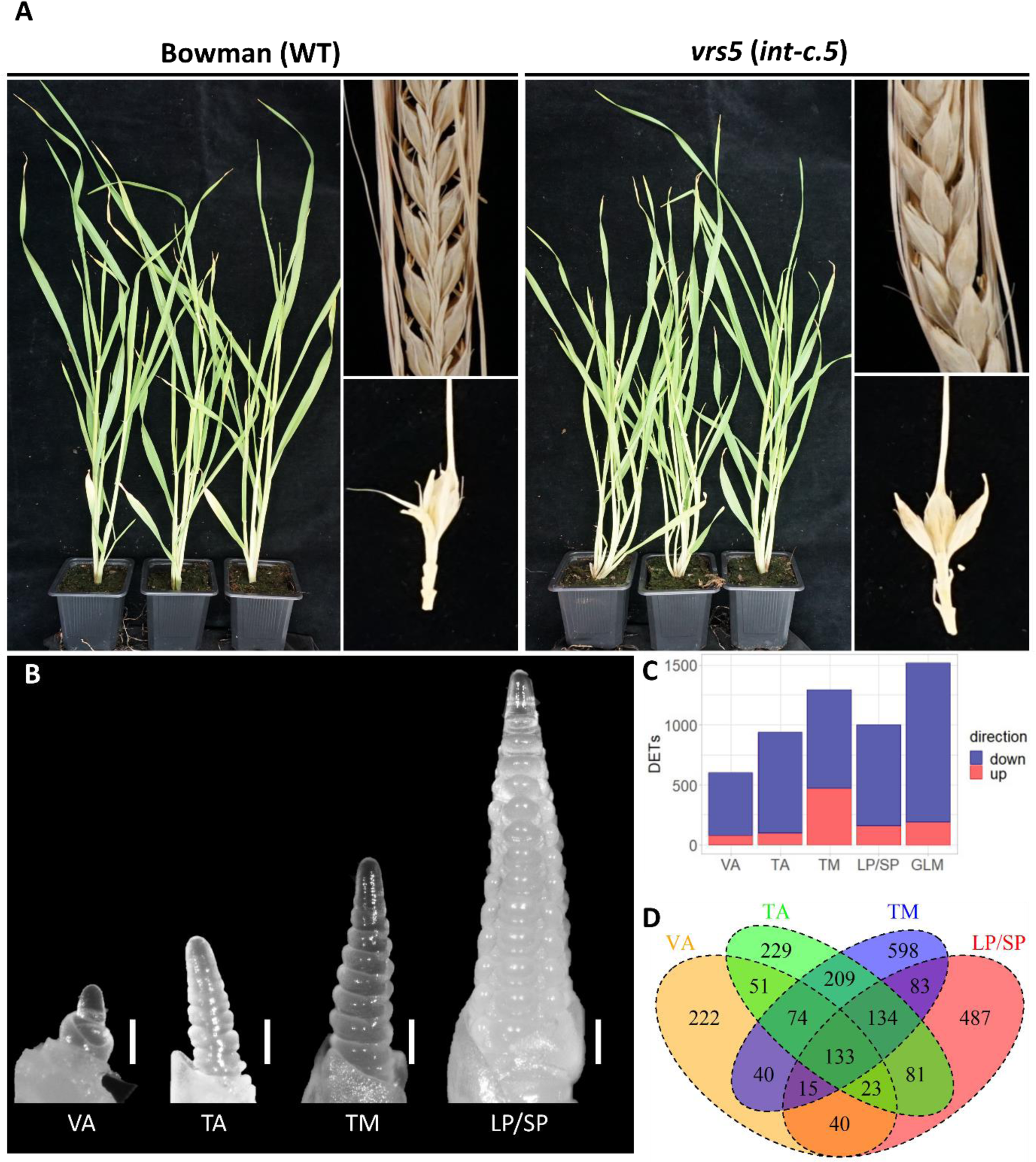
VRS5 regulates both tiller number and lateral spikelet development in barley. A) Four-week-old plants, dried spike and spikelet triplet of WT (Bowman) and vrs5 (int-c.5) respectively. The lateral spikelets of WT have not developed, whereas the lateral spikelets of vrs5 have developed into grains. B) binocular images of developing inflorescence meristems representing the four stages used for transcriptome data, namely: vegetative apex (VA), transition apex (TA), triple mound (TM), and lemma and stamen primordia (LP/SP). The white bar represents a length of 200 µm. C) Bar graph showing number of differentially expressed transcripts (DETs) per developmental stage including the direction of differential expression in vrs5 compared to Bowman. D) Venn-diagram showing overlap of DETs at different developmental stages.

From the generated transcriptomes, principal component analysis (PCA) of all expressed transcripts showed that developmental stage (PC1) and the genotype (PC2) contribute the most to the observed variation (**Figure S3**). To gain further insight into the differentially expressed genes between *vrs5* and Bowman, we compared the transcriptomes in two different ways, namely at each developmental stage specifically (VA, TA, TM, and LP/SP), and over development using a generalized linear model (GLM) (**Table S2-S6**). When comparing differentially expressed transcripts (DETs) at each developmental stage, most DETs (1286) were found at TM, whereas at VA the least DETs (598) were found (**Figure 1C**). Comparing the overlap in DETs in the different stages revealed that 133 transcripts were different between *vrs5* and Bowman in all developmental stages (**Figure 1D**). Of the total amount of DETs found, the majority (∼75%) was downregulated in *vrs5* compared to Bowman, indicating that VRS5 mainly acts as a transcriptional activator (**Figure 1**).

### Transcriptional profiling of genes differentially regulated in *vrs5*

To gain a deeper understanding of differential gene expression between the wild type and *vrs5* mutant over development, a co-expression clustering was performed using 2,872 DETs that were differentially regulated in one, or more stages or independent of development. This yielded 39 clusters, which were subdivided into three main groups based on their expression pattern (**Figure 2A, Figure S4, Table S9**): Group 1 represents genes that are highly expressed at VA and TA, and low expressed at TM and LP/SP developmental stages; Group 2 represents genes that are lowly expressed at VA and TA, and increase in expression during TM and LP/SP stages; Group 3 represents genes that have a similar expression over the developmental stages (**Figure 2A**). Group 1 has clearly the most DETs, from which the majority, are in group 1A (1,776, ∼62%), which consists of DETs that go down in expression over time and have lower expression in *vrs5* (**Figure 2A**).

**Figure 2:**
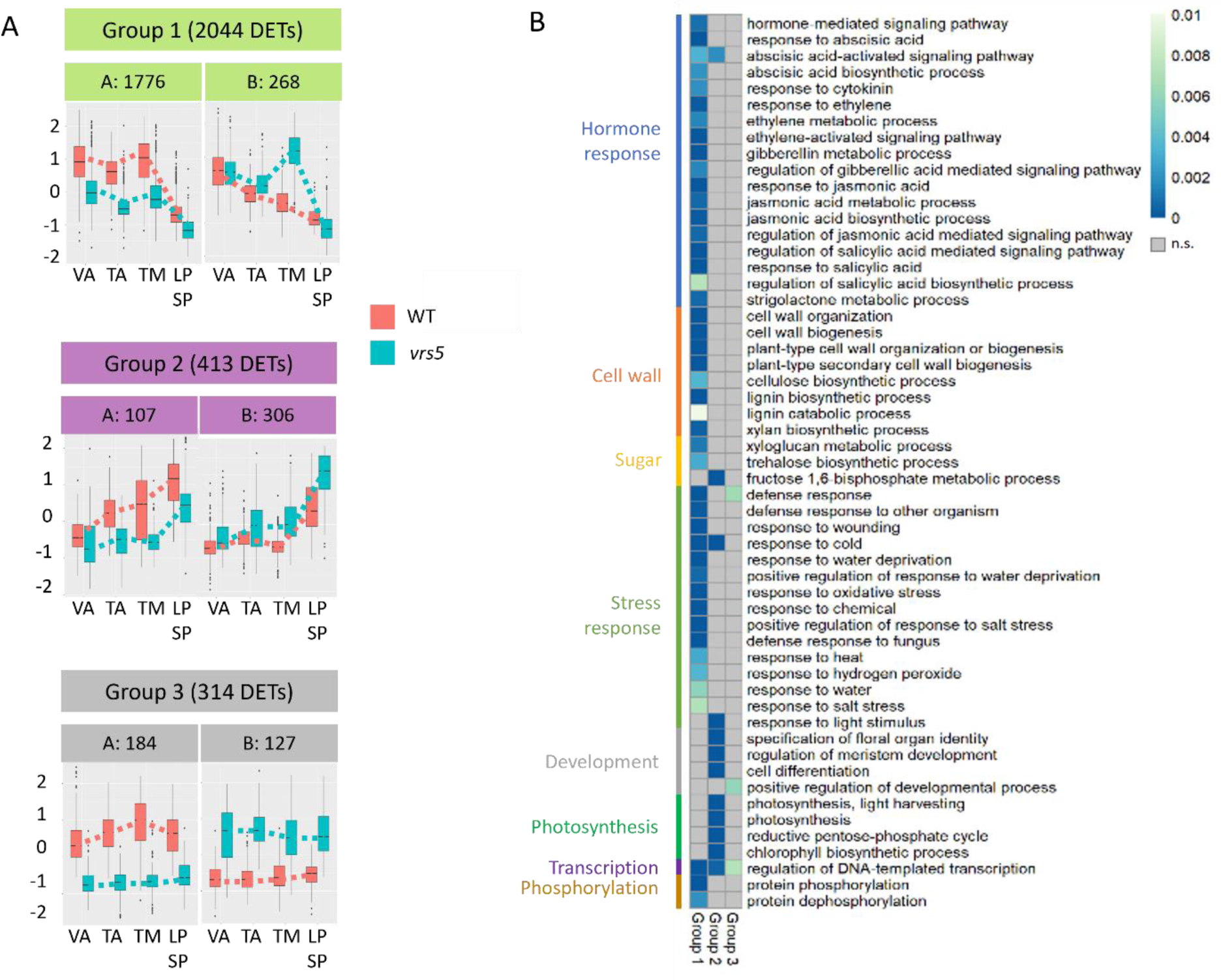
Transcriptional profiling of VRS5 shows changes in gene expression during development. A) Boxplots of z-scores of DETs in three main co-expression cluster groups. Boxplots are colored red for Bowman and cyan for vrs5. B) Enriched GO-terms for Biological Processes per cluster group where the more blue means a lower p-value. A grey color means that the GO-term is not enriched in that cluster group (p > 0.01) or the number of DETs belonging to that GO-term are lower than three.

To gain insight in the downstream processes VRS5 regulates during development, GO-term enrichment was performed on the expression groups. Genes that are highly expressed during VA and TA (Group 1), and differentially regulated in *vrs5* when compared to wild-type lines are mostly enriched for GO-terms related to protein phosphorylation and phytohormone response and signaling (**Figure 2B**). DETs in group 1, annotated with these GO terms include several Jasmonate ZIM-domain containing proteins (JAZ), involved in jasmonic acid (JA) signaling (*HORVU.MOREX.r3.2HG0163880, HORVU.MOREX.r3.5HG0480600, HORVU.MOREX.r3.4HG0405240*, and *HORVU.MOREX.r3.2HG0125430*). Genes involved in abscisic acid (ABA) biosynthesis and signaling are also enriched in group 1, like HORVU.MOREX.r3.4HG0383820, a 9-cis-epoxycarotenoid dioxygenase (NCED) involved in ABA biosynthesis, and HORVU.MOREX.r3.5HG0508150, an ABA receptor. Orthologs of these genes are also differentially expressed in maize *tb1* mutants compared to WT (Dong et al., 2019), suggesting a conserved regulation by the transcription factor proteins encoded by the orthologous genes *ZmTB1* and *VRS5*. Next to GO-terms involved in phytohormone signaling, GO-terms involved in stress response are highly enriched early in development.

Genes in group 2 are highly expressed at the TM and LP/SP stages when compared to the VA and TA. GO-overrepresentation analysis of DETs between Bowman and *vrs5* lines showed that in this group genes that are involved in “regulation of meristem development”, “plant organ development”, and “cell differentiation” are overrepresented (**Figure 2B**). This may reflect the subsequent development of lateral florets in *vrs5* in contrast to their repression in WT. In line with this, we also identified floral homeotic genes amongst the differentially regulated genes including *HORVU.MOREX.r3.1HG0065060*, homologous to *Arabidopsis PISTILLATA*; and all identified E-class MIKCc MADS box genes in barley (*HORVU.MOREX.r3.4HG0396400*, *HORVU.MOREX.r3.7HG0654930, HORVU.MOREX.r3.5HG0511250, HORVU.MOREX.r3.7HG0684020,* and *HORVU.MOREX.r3.5HG0494190)* (Kuijer et al., 2021). Also other MIKCc MADS box genes were differentially expressed in *vrs5*, including *HORVU.MOREX.r3.6HG0604360*, orthologous to rice *MADS6,* and wheat *AGL6* (Kuijer et al., 2021). In wheat, *agl6* mutants are infertile due to defective pistils. Moreover, *TaAGL6* was shown to act upstream of several floral homeotic genes (Kong et al., 2022).

Our results show that VRS5 during early, tiller developmental, stages is targeting amongst others, phytohormone and stress signaling associated genes. In contrast, during inflorescence development mainly genes involved in meristem development and floral organ identity are differentially regulated in the absence of VRS5. Together, our results suggest that VRS5 regulates different genes and processes depending on the specific developmental stage.

### Genome wide identification of the *bona fide* VRS5 targets

The transcriptome profiling showed that there are differences in the differentially regulated genes at early, tiller versus inflorescence developmental stages. As a transcription factor, VRS5 is expected to regulate the expression of downstream target genes by directly binding to the (distal) promoter region of these genes. As key regulator, VRS5 may only bind to a limited number of downstream genes to trigger a wider transcriptional response. Alternatively, VRS5 may directly activate the majority of the genes identified as DET. To gain more insight into which genomic regions are bound by VRS5, and which of these are within a promoter region of a gene, we performed a DNA-affinity purification followed by genome wide sequencing (DAP-Seq) (Bartlett et al., 2017). To construct our DAP-Seq libraries, we used DNA isolated from the barley apical meristem. VRS5 fused to a FLAG-tag was used as bait, while GFP, which is expected not to bind DNA, fused to a FLAG-tag was used as negative control. Regions in the genome enriched for VRS5-binding compared to GFP are called peaks and considered genomic VRS5 binding sites. Only the peaks with an adjusted p-value smaller than 1^-10^ were kept, which resulted in 99,567 peaks **(Table S7).** The sequences of all the peaks were analyzed for de-novo enriched motifs. The most frequently abundant motif was GGGNCCCAC, found in 72% of the peaks **(Figure 2A).** This motif was distributed closely around the peak summit **(Figure 2B),** indicating the GGGNCCCAC motif is the primary DNA motif bound by VRS5. This motif resembles a consensus class II TCP-binding motif (GGGNCCAC), and the consensus binding motifs found for homologous TCP transcription factors in *Arabidopsis* and maize (Dong et al., 2019; Gonzalez-Grandio et al., 2017; Kosugi and Ohashi, 2002; van Es et al., 2023). This supports the association of identified peaks as likely genomic VRS5 binding sites. To assign peaks to putative target genes, we selected a region of 15kb up and downstream of all predicted barley open reading frames. Within these genomic regions, we found 21% of all identified VRS5-DAP-seq peaks. The other peaks were located outside this region and classified therefore as “Distal intergenic”. Only a few binding sites, 0.1% of the total number of peaks, were identified in intronic regions (**Table S8**). To further understand where the peaks are located in the promoter and downstream region of the gene, we generated a frequency plot of all peaks that were found in genes that are expressed in at least one of the VA, TA, TM, or LP/SP stages. Results showed that peaks in expressed genes, were mostly found to bind the promoter or terminator rather than the gene body (**Figure 3C**). By combining the VRS5 binding peaks with the DETs found in our expression analysis between *vrs5* and WT, we identified putative direct downstream targets of VRS5 (*bona fide* VRS5 targets). To evaluate if these are equally distributed over the three expression groups identified in the co-expression clustering, we compared the percentage of genes that were differentially regulated in the transcriptional profiling, and of which the promoter and/or terminator was bound by VRS5 within the DAP-Seq experiment. Results showed that in all three expression groups, the percentage of DETs with a VRS5-DAP-Seq peak was similar (∼20-25%). This suggests that VRS5 is directly regulating transcription of its targets throughout the developmental stages tested.

**Figure 3:**
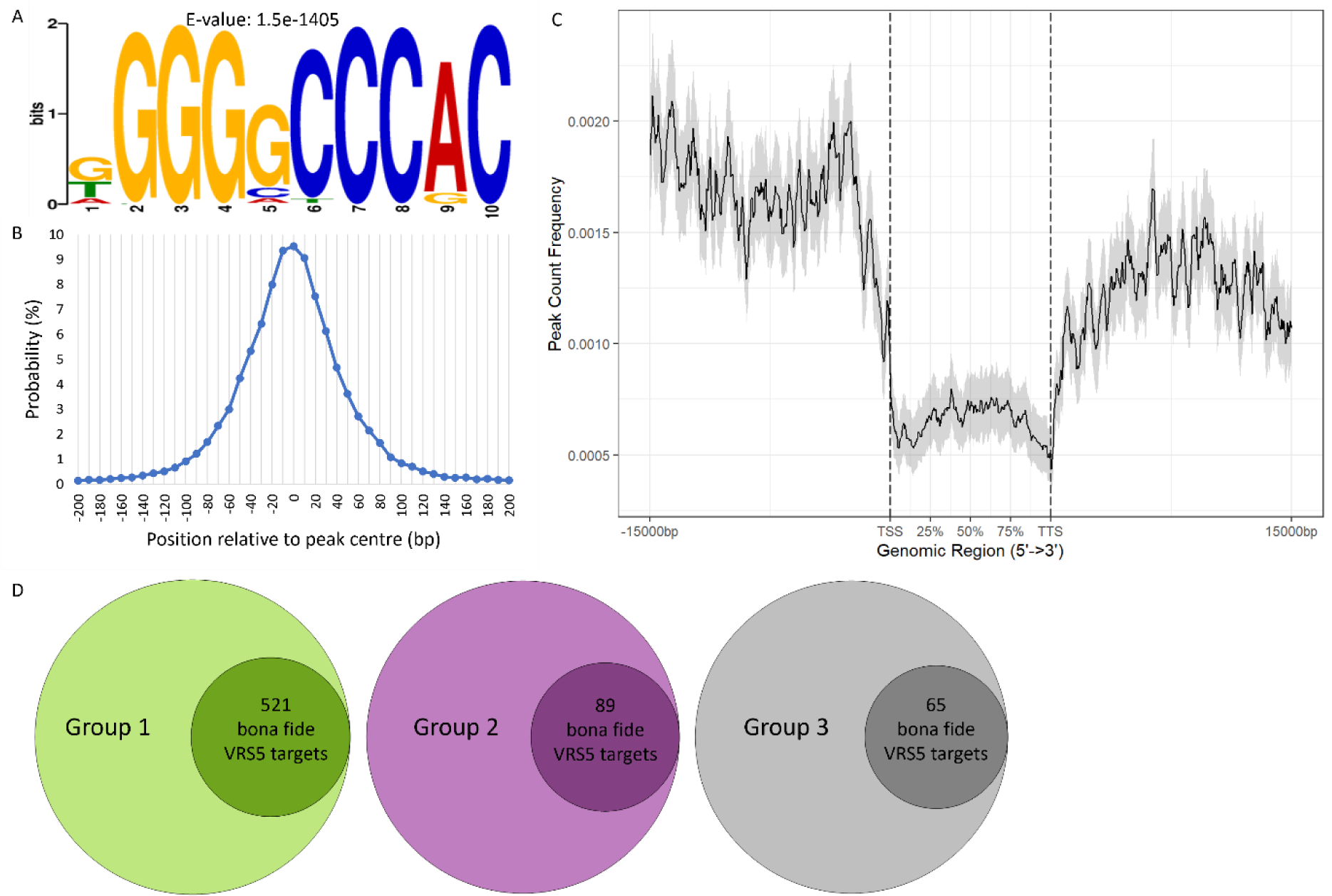
DAP-Seq of VRS5 on barley genomic DNA library. A) Enriched sequence logo in the VRS5 bound DAP-seq peaks, identified using STREME. B) The position of the motif shown in A relative to the center of the VRS5 DAP-seq peak as a probability percentage of bins of 10 bp. C) Frequency distribution plot of the VRS5 peaks 15kb upstream of the TSS and 15 kb downstream of TTS along expressed genes in the developmental stages sampled. D) Differentially regulated transcripts (DETs) per group that have a VRS5 binding peak within 15kb of their open reading frame in the DAP-Seq experiment.

### VRS5 binds to the promoter of genes involved in inflorescence development

To further explore the role of VRS5 in regulating genes involved in inflorescence development, we focus on the *bona fide* VRS5 targets identified in group 2, which comprises DETs that are low at VA and TA and high at the LP/SP stages. To extend the analysis on *bona fide* VRS5 targets with the stamen primordia (SP) and awn primordia (AP) stages, we re-analyzed available RNA-Seq (van Esse et al., 2017) and included DETs identified (**Table S10**). We identified several known genes in inflorescence and flower development, like a C2H2 zinc finger (*HORVU.MOREX.r3.2HG0170820*), which is identified as candidate for the long glume mutant (*lgm1)* in barley (Z. Zhang et al., 2024). Its ortholog in wheat is causing awn suppression (DeWitt et al., 2020; Huang et al., 2020). Other targets known in inflorescence and flower development include several MIKC-type MADS-box genes, which have a conserved role in the specification of floral organ identity (Kuijer et al., 2021). Amongst these are orthologs of floral organ development genes such as *PISTILLATA* (*HORVU.MOREX.r3.1HG0065060)*, *MADS1* (*HORVU.MOREX.r3.4HG0396400), AGL6 (HORVU.MOREX.r3.6HG0604360)*, and *SOC1-like* (*HORVU.MOREX.r3.1HG0054220*), that also contained a DAP-Seq peak.

*CENTRORADIALIS (HvCEN; HORVU.MOREX.r3.2HG0166090),* is a gene involved in the regulation of meristem development, and downregulated in *vrs5* throughout development compared to Bowman (**Figure 3A**). Next to that, a VRS5 DAP-peak was found ∼7 kb upstream of the coding sequence of *CENTRORADIALIS*. HvCEN acts as a floral repressor and mutants show early flowering, shorter spikes and less tillers at maturity (Bi et al., 2019; Comadran et al., 2012). To confirm the DAP-Seq results, we performed an EMSA assay using probes of DNA sequences extracted from the DAP-peak that contained the VRS5 binding site GGGNCCCAC. We used the DAP-peaks identified within the *HvCEN, PISTILLATA*, and *SOC1-like* promoters. The peak in the promoter of *HvCEN* contained two putative VRS5 binding sites, while in the *PISTILLATA* and *SOC1-like* promoter regions, one was found. Our results showed that VRS5 can indeed bind to the core VRS5 binding motif GGGNCCCAC in these promoters (**Figure 4B**).

**Figure 4:**
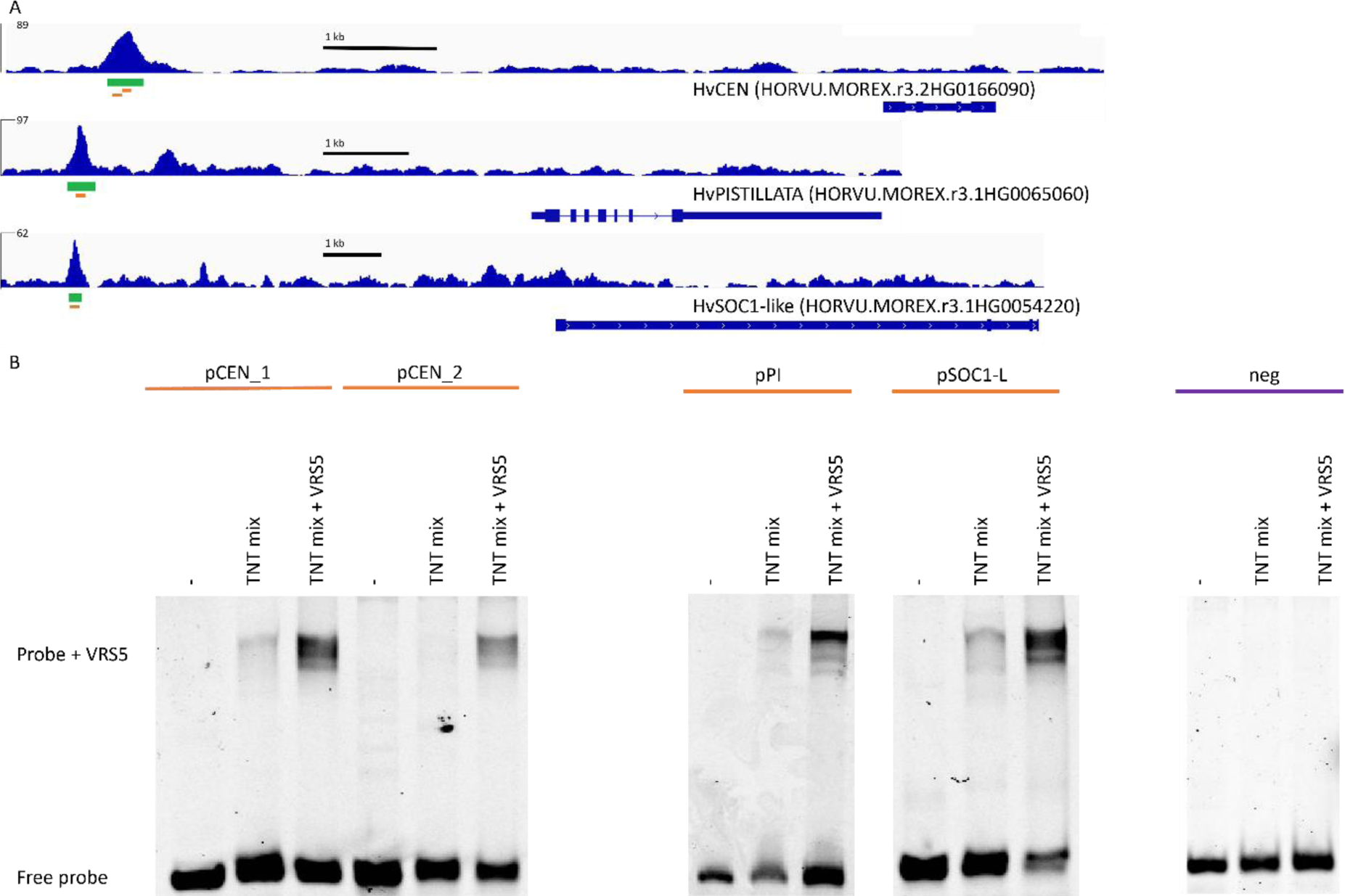
VRS5 binds to the promoter of genes involved in inflorescence development. A) VRS5 DAP-seq peaks identified within the PISTILLATA, SOC1-like and HvCEN promoter. The green bars represent the peak significant compared to the control. The orange bars mark the VRS5 binding sites within the peaks. B) Confirmation of the VRS5 binding to DAP-Seq peaks using EMSA assays on the VRS5 binding sites identified within the promoter of HvCEN (pCEN), PISTILLATA (pPI), and SOC1-like (pSOC11-L). The HvCEN promoter contained two putative VRS5 binding sites within the DAP peak, which are indicated as pCEN_1 and pCEN_2. As negative control a probe that does not contain a TCP binding motif outside of a DAP-seq peak was taken.

### DAP-Seq and RNA-Seq reveal a conserved mode of action of VRS5

In the dicot model plant *Arabidopsis* BRC1, a close homolog of *VRS5*, directly binds to the promoter of HD-Zip genes HB21, HB40, and HB53 to regulate their expression (Gonzalez-Grandio et al., 2017; van Es et al., 2023). The regulation of HD-Zip genes is conserved in maize, where ZmTB1 directly binds to the promoter of *GRASSY TILLERS* (*ZmGT1*). Given the conserved role of TB1 and BRC1 in regulation of HD-ZIP transcription factors, we first explored if amongst our *bona fide* targets HD-ZIP TF were present. Interestingly, the row-type related gene *VRS1* (*HORVU.MOREX.r3.2HG0184740*) was amongst the *bona fide* downstream targets of VRS5 (Figure 5, Table S10). *VRS1* encodes a HD-ZIP protein that is closely related to *ZmGT1* and the *Arabidopsis* HD-Zip genes. VRS1 is well known for its role in lateral spikelet development. Plants that do not have functional VRS1 exhibit a six-rowed spike architecture (Komatsuda et al., 2007, Liller et al., 2015). Our targeted mutagenesis using CRISPR corroborated the crucial role of *VRS1* in inhibiting the outgrowth of lateral florets (**Figure 5A, B**). The VRS5 binding peak in our DAP-seq experiment was found around 7kb upstream of the *VRS1* translational start site. Sequence analysis of the region of this DAP-peak showed that there are three putative VRS5 binding sites within the center of this peak (**Figure 5C**). To verify if VRS5 was indeed capable of binding to the core VRS5 binding motifs found in this region, we performed an EMSA targeting all three putative VRS5 binding sites separately. This assay revealed that VRS5 was capable of binding to all three putative binding sites (**Figure 5D**), providing evidence that VRS5 can directly bind to the *VRS1* promoter to regulate *VRS1* expression.

**Figure 5:**
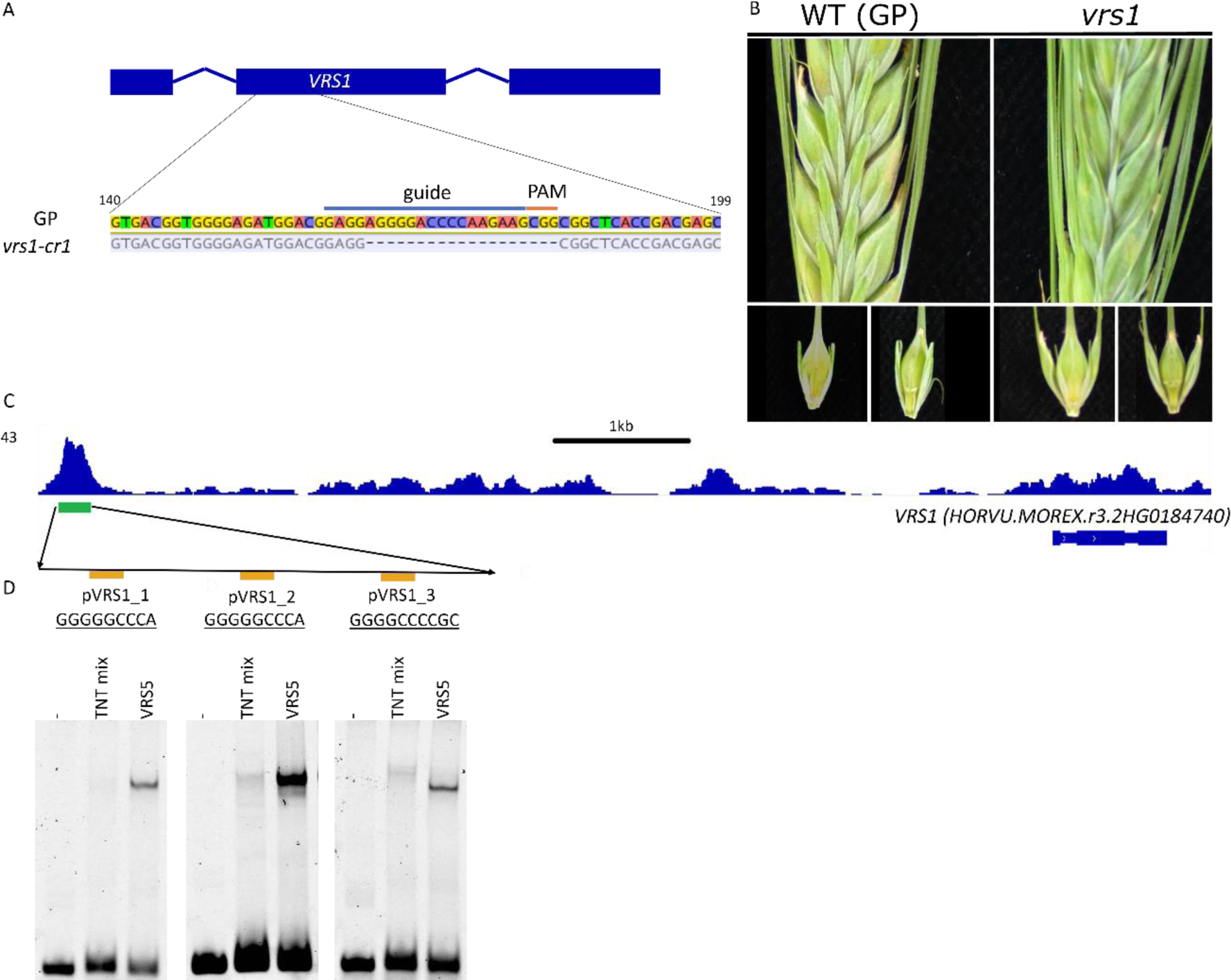
VRS5 binds to the promoter of VULGARE ROWTYPE SIX 1 (VRS1) to regulate its expression. A) Gene editing of vrs1 with targeted mutagenesis (CRISPR). B) Phenotype of gene edited vrs1 mutant in the genetic background of GOLDEN PROMISE (GP) compared to the GP parental line. VRS1 lines have a six-rowed spike architecture. C) VRS5 binding region identified with DAP-Seq within the promoter of VRS1. There are three putative VRS5 binding sites within the VRS1 DAP-Seq peak. D) EMSA confirms the DAP-Seq data that demonstrated binding of VRS5 to the three VRS5 binding sites within the VRS1 promoter (pVRS1).

**Figure 6:**
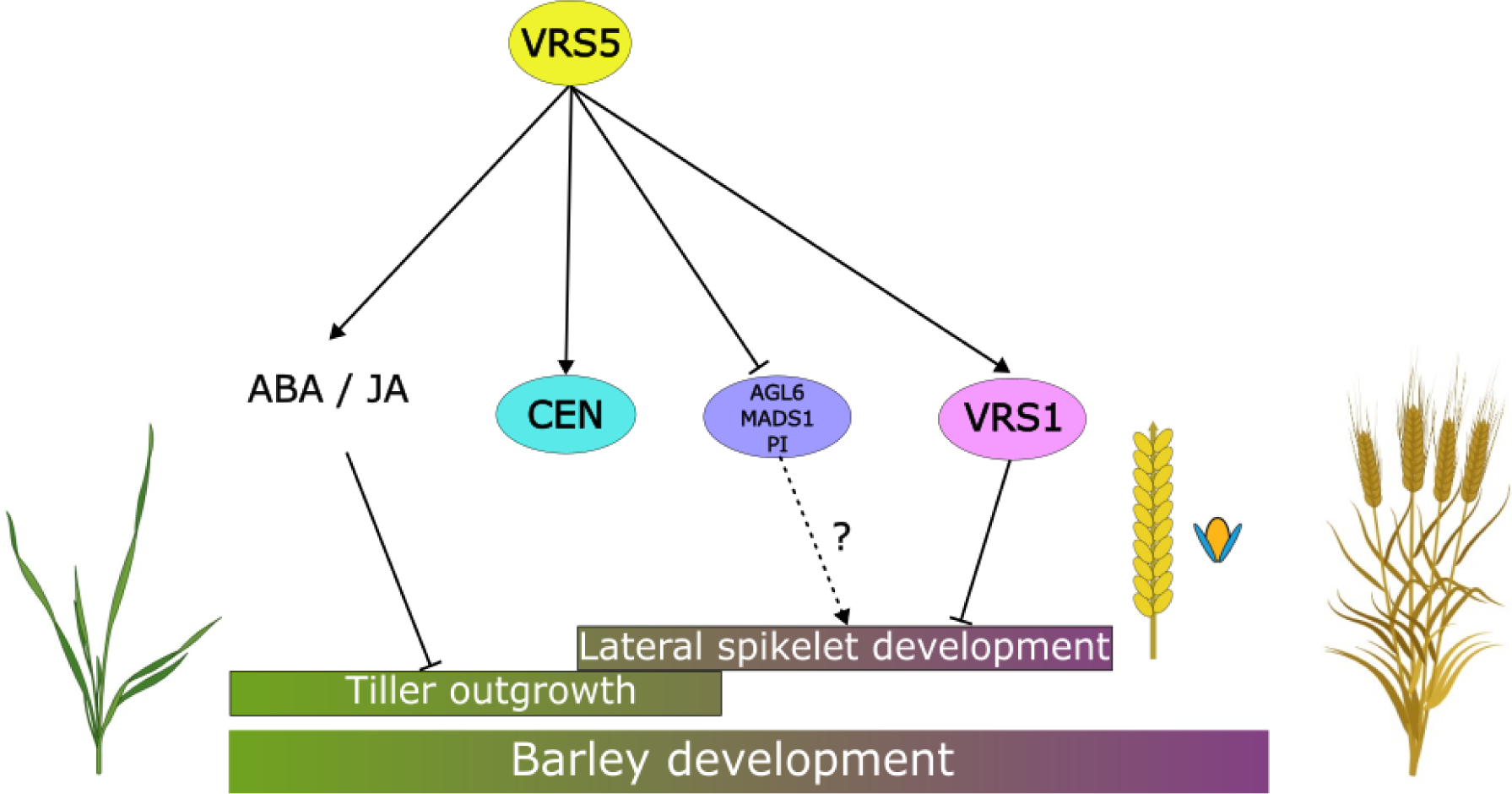
VRS5 network over development. VRS5 inhibits tiller outgrowth via ABA/JA early in development. Later in development, VRS5 inhibits lateral spikelet development directly via VRS1. VRS5 also inhibits expression of AGL6, MADS1, and PISTILLATA (PI) directly, which might play a role in lateral spikelet development. Additionally, VRS5 induces the expression of HvCEN.

Aside from *VRS1*, our transcriptional profiling combined with GO annotation showed that genes which are highly expressed at VA and TA and low at the LP/SP stages (Group 1) are overrepresented for ABA and JA related genes. Previously, it was demonstrated in maize that ABA and JA related genes are among the putative downstream targets of ZmTB1 in tiller development (Dong et al., 2019). To further explore which *bona fide* VRS5 targets are conserved in maize, we used ZmTB1 targeted ABA and JA related genes and identified their orthologous genes in barley. All of the identified barley orthologs were downregulated in *vrs5* compared to WT and most were grouped in cluster group 1A, based on their expression pattern (**Table S11**). This suggests that these genes are mostly import early in development, when the tillers are formed. We found that indeed ZmTB1 and VRS5 share overlapping *bona fide* ABA and JA-related targets: AP2/B3 transcription factor *RELATED TO ABI3/VP 2 (RAV2) (HORVU.MOREX.r3.3HG0281730)*; several lipoxygenases, including *TASSELSEED1* (*HORVU.MOREX.r3.2HG0171680)*; and *JAZs* (**Table S11**). This shows that the regulation of ABA and JA related genes is a conserved function between barley VRS5 and the maize ortholog TB1.

## Discussion

VRS5 is an important domestication gene in barley that plays a crucial role in tillering and row-type determination (Liller et al., 2015; Ramsay et al., 2011; Zwirek et al., 2019). Using a combination of transcriptomics and localizing genome-wide binding sites of VRS5, we identified its putative direct downstream targets. Different from ChIP-sequencing, which is a more *in-vivo* method to find TFBS and where inaccessible DNA cannot be bound, DAP-sequencing is a method where naked DNA is used, so chromatin context is not kept (Bartlett et al., 2017; Hajheidari and Huang, 2022). Nevertheless, despite the lack of chromatin context in DAP-seq, the overlap between DAP-seq and ChIP-seq peaks is generally high (O’Malley et al., 2016). Of the DAP peaks found for VRS5, a significant part was found distal intergenic. A recent study performed DAP-sequencing of a large number of transcription factors in *Triticum urartu,* that has a genome size comparable to barley. They found for the majority of transcription factors that more than 50% of the DAP-seq peaks were located more than 10kb from the gene body (Zhang et al., 2021). Similarly, in strawberry DAP-Seq of a TCP transcriptional regulator revealed also presence of the majority of peaks in the distal intergenic region (Chen et al., 2024). Together, this indicates that for DAP-sequencing on a large genome such as barley, a large portion of distal intergenic peaks can be expected. As an in vitro technique, DAP-sequencing might yield binding sites that would never be accessible under *in-vivo* circumstances. In part, this could result in a higher number of distal intergenic DAP-peaks (Maher et al., 2018). To what extend distal intergenic DAP-peaks in general, and for VRS5 in this study, represent biological relevant long distal regulatory elements still needs to be elucidated. In barley, the most accessible chromatin regions (ACRs) are found between 10 and 100 kb from the nearest gene (Lu et al., 2019). Recent studies using STARR-Seq indicate that about ∼37% of the enhancers had potential target genes at a distance of 10kb, suggesting that only a minority of the genes in barley is regulated by adjacent enhancers (Zhou et al., 2024). In our study, we focused on DAP-peaks found within 15 kb of their nearest gene whilst a significant portion of the DAP-Seq peaks was found in distal-intergenic regions. Given the observation that in barley, enhancers may lie up to 100 kb (Zhou et al., 2024) from a target gene, biological relevance of other DAP-peaks beyond our window cannot be excluded. Knowing the ACRs in tissues where VRS5 is expressed and under different environmental circumstances could give a better understanding and biological relevance to these other peaks. Taken together, to truly place distal-intergenic peaks into a biological context, a more in-depth study of the 3-D genome makeup at high resolution is imperative.

Nevertheless, in our analysis we identified >700 *bona fide* VRS5 targets and reveals an overlap in ABA and JA-related targets in maize and barley. This suggests that the network of tiller development via ABA and JA signaling is partially conserved between barley and maize. In the model plant *Arabidopsis,* ABA related genes are directly targeted by the class II TCP transcription factor BRC1 to regulate axillary branching (Gonzalez-Grandio et al., 2017; van Es et al., 2023). This implies that the regulation of ABA signaling by BRC1/TB1 proteins is evolutionary ancient and precede the divergence of monocotyledonous and dicotyledonous plants (van Es et al., 2023). Interestingly, some of these genes found as *bona fide* VRS5 target may have a role in inflorescence development, from which the most known gene is *VRS1*. Intriguingly, VRS1 and VRS5 alleles have been co-selected for increased lateral grain size in six-rowed cultivars (Ramsay 2011), and *vrs5 vrs1* double mutants have an additive effect when compared to the respective single mutants (Zwirek et al., 2019). In addition, *vrs5* mutants have an intermediate phenotype, whereas *vrs1* lines have a complete six-rowed phenotype (Liller et al., 2015; Zwirek et al., 2019). Combined, this suggest that there are other activators of *VRS1* that may act redundant, or in parallel to VRS5.

Aside from VRS1, our analysis also revealed other genes involved in inflorescence development amongst the *bona fide* VRS5 targets. For example, we identified a ortholog of TASSELSEED 1 as *bona fide* target of VRS5. In maize, *TASSELSEED1* is a lipoxygenase involved in JA biosynthesis and known for its important role in sex-determination of maize florets by repressing carpel development (Acosta et al., 2009). Furthermore, in our data we observed an upregulation of genes involved in flowering and floral organ identity, such as MIKC-type MADS-box genes, in *vrs5* compared to WT. MIKC-type MADS-box genes, which include the ABCDE-genes that are known for regulating floral homeotic patterning, are mostly upregulated during inflorescence development in barley (Digel et al., 2015; Kuijer et al., 2021; Liu et al., 2020). Our observation that floral homeotic genes are upregulated in *vrs5* when compared to Bowman may reflect an increased expression of these genes in lateral spikelet primordia in *vrs5*. However, the presence of a VRS5 DAP-peak within the promoter region of some of these MIKC-type MADS-box genes suggests a more direct effect of VRS5 on the regulation of these floral homeotic genes. Intriguingly, *HvMADS1*, which in barley regulates lemma and awn development and prevents inflorescence branching under high ambient temperatures (Li et al., 2021; Y. Zhang et al., 2024) is among the *bona fide* VRS5 targets. Moreover, an ortholog of AGL6 was identified as *bona fide* VRS5 target that is significantly upregulated in the *vrs5* mutant when compared to WT. In wheat, *agl6* mutants are infertile due to defective pistils and in this species, AGL6 acts upstream of several other floral homeotic genes (Kong et al., 2022). These results suggest that VRS5 may also inhibit lateral spikelet development in two-rowed cultivars by directly reducing the expression of other genes involved in floral organ development. In concurrence with this hypothesis, row-type genes prevent the development of carpels in the two-rowed spike architecture (Zwirek et al., 2019), while loss of function mutants of *vrs5* and other row-type genes have an increased lateral floret fertility.

Aside from the floral homeotic genes, *HvCEN* is also identified amongst the *bona fide* targets VRS5. HvCEN acts antagonistically to FLOWERING LOCUS-T (FT) in regulating floral transition (Bi et al., 2019; Colleoni et al., 2024). In wheat and *Arabidopsis*, TaTB1-D and AtBRC1 respectively, are shown to interact with the FT1 ortholog of their species (Dixon et al., 2020; Niwa et al., 2013). To what extent binding of VRS5 to the HvCEN promoter impacts on floral transition, flowering or meristem development remains to be elucidated. However, our results do suggest a role of VRS5 in the direct regulation of *PEBP* genes through binding to the promoter to activate the expression aside from the previously described regulation through protein-protein interaction.

In conclusion, with this study, we showed that barley VRS5 binds a class II TCP DNA motif GGGNCCCAC and acts mainly as a transcriptional activator. *Bona fide* VRS5 target genes include genes involved in ABA-signaling, which seems to be a conserved role for TB1-like proteins in both monocots and dicots to regulate tillering or axillary branching. We also revealed a role for VRS5 in regulation of floral homeotic and meristem identity genes. Our study exemplifies that it is essential to not only identify the molecular networks in model plants like *Arabidopsis*, but also in key crops such as barley to reveal conservation, but more important to identify divergence explaining the species-specific architecture. The study presented here offers a solid and crucial starting point for mutant analyses of a selection of *bona fide* targets to gain a deeper understanding of the VRS5 network and to enlighten its mode of action in shaping barley development.

## Acknowledgements and funding

We acknowledge Amy Raphaella for generating constructs used in the EMSA assays during her BSc thesis project. This work was financially supported by a NWO-VIDI grant to GWvE (project number 18366), the UKRI-BBSRC-ISP (grant no. BB/P012574/1 and BBS/E/J/000PR9795 to MJM), Gatsby Charitable Foundation (MJM), and United States Department of Agriculture-Agricultural Research Service CRIS #5062-21220-025-000D (MJM).

## Author contributions

T.W. designed and performed research, generated and analyzed DAP-Seq and RNA-Seq data and performed the EMSA assays. P.C. designed research and contributed to the generation of the RNA-Seq data. R.I. and G.W.v.E designed research. M.J.M. provided the *vrs1* mutants, and critical discussions. R.I., G.W.v.E, P.H., K.O., and R.C.S. participated in the design of the experiments and critical discussions. T.W., G.W.v.E. co-wrote the manuscript with contributions from all co-authors.

## References

Acosta, I.F., Laparra, H., Romero, S.P., Schmelz, E., Hamberg, M., Mottinger, J.P., Moreno, M.A., Dellaporta, S.L., 2009. Tasselseed1 Is a Lipoxygenase Affecting Jasmonic Acid Signaling in Sex Determination of Maize. Science 323, 262–265.

Bailey, T.L., 2021. STREME: accurate and versatile sequence motif discovery. Bioinformatics 37, 2834–2840. 10.1093/bioinformatics/btab203

Bailey, T.L., Johnson, J., Grant, C.E., Noble, W.S., 2015. The MEME Suite. Nucleic Acids Res. 43, W39–W49. 10.1093/nar/gkv416

Barrett, T., Wilhite, S.E., Ledoux, P., Evangelista, C., Kim, I.F., Tomashevsky, M., Marshall, K.A., Phillippy, K.H., Sherman, P.M., Holko, M., Yefanov, A., Lee, H., Zhang, N., Robertson, C.L., Serova, N., Davis, S., Soboleva, A., 2013. NCBI GEO: archive for functional genomics data sets—update. Nucleic Acids Res. 41, D991–D995. 10.1093/nar/gks1193

Bartlett, A., O’Malley, R.C., Huang, S.S.C., Galli, M., Nery, J.R., Gallavotti, A., Ecker, J.R., 2017. Mapping genome-wide transcription-factor binding sites using DAP-seq. Nat. Protoc. 12, 1659–1672. 10.1038/nprot.2017.055

Bi, X., Esse, W.V., Mulki, M.A., Kirschner, G., Zhong, J., Simon, R., Korff, M.V., 2019. Centroradialis interacts with flowering locus t-like genes to control floret development and grain number. Plant Physiol. 180, 1013–1030. 10.1104/pp.18.01454

Bolger, A.M., Lohse, M., Usadel, B., 2014. Trimmomatic: A flexible trimmer for Illumina sequence data. Bioinformatics 30, 2114–2120. 10.1093/bioinformatics/btu170

Chen, X., Gao, J., Shen, Y., 2024. Abscisic acid controls sugar accumulation essential to strawberry fruit ripening via the FaRIPK1-FaTCP7-FaSTP13/FaSPT module. Plant J. n/a. 10.1111/tpj.16862

Clark, R.M., Linton, E., Messing, J., Doebley, J.F., 2004. Pattern of diversity in the genomic region near the maize domestication gene tb1. Proc. Natl. Acad. Sci. 101, 700–707. 10.1073/pnas.2237049100

Colleoni, P.E., van Es, S.W., Winkelmolen, T., Immink, R.G.H., van Esse, G.W., 2024. Flowering time genes branching out. J. Exp. Bot. erae112. 10.1093/jxb/erae112

Comadran, J., Kilian, B., Russell, J., Ramsay, L., Stein, N., Ganal, M., Shaw, P., Bayer, M., Thomas, W., Marshall, D., Hedley, P., Tondelli, A., Pecchioni, N., Francia, E., Korzun, V., Walther, A., Waugh, R., 2012. Natural variation in a homolog of Antirrhinum CENTRORADIALIS contributed to spring growth habit and environmental adaptation in cultivated barley. Nat. Genet. 44, 1388–1391. 10.1038/ng.2447

Dang, Y., Jia, G., Choi, J., Ma, H., Anaya, E., Ye, C., Shankar, P., Wu, H., 2015. Optimizing sgRNA structure to improve CRISPR-Cas9 knockout efficiency. Genome Biol. 16, 280. 10.1186/s13059-015-0846-3

DeWitt, N., Guedira, M., Lauer, E., Sarinelli, M., Tyagi, P., Fu, D., Hao, Q., Murphy, J.P., Marshall, D., Akhunova, A., Jordan, K., Akhunov, E., Brown-Guedira, G., 2020. Sequence-based mapping identifies a candidate transcription repressor underlying awn suppression at the B1 locus in wheat. New Phytol. 225, 326–339. 10.1111/nph.16152

Digel, B., Pankin, A., von Korff, M., 2015. Global Transcriptome Profiling of Developing Leaf and Shoot Apices Reveals Distinct Genetic and Environmental Control of Floral Transition and Inflorescence Development in Barley. Plant Cell 27, 2318–2334. 10.1105/TPC.15.00203

Dixon, L.E., Greenwood, J.R., Bencivenga, S., Zhang, P., Cockram, J., Mellers, G., Ramm, K., Cavanagh, C., Swain, S.M., Boden, S.A., 2018. TEOSINTE BRANCHED1 regulates inflorescence architecture and development in bread wheat (Triticum aestivum). Plant Cell 30, 563–581. 10.1105/tpc.17.00961

Dixon, L.E., Pasquariello, M., Boden, S.A., 2020. TEOSINTE BRANCHED1 regulates height and stem internode length in bread wheat. J. Exp. Bot. 71, 4742–4750. 10.1093/jxb/eraa252

Dixon, L.E., Van Esse, W., Hirsz, D., Willemsen, V., McKim, S.M., 2022. Cereal Architecture and Its Manipulation. Annu. Plant Rev. Online 5, 1–54. 10.1002/9781119312994.apr0648

Doebley, J., Stec, A., Gustus, C., 1995. teosinte branched1 and the origin of maize: evidence for epistasis and the evolution of dominance. Genetics 141, 333–346. 10.1093/genetics/141.1.333

Doebley, J.F., Gaut, B.S., Smith, B.D., 2006. The Molecular Genetics of Crop Domestication. Cell 127, 1309–1321. 10.1016/j.cell.2006.12.006

Dong, Z., Xiao, Y., Govindarajulu, R., Feil, R., Siddoway, M.L., Nielsen, T., Lunn, J.E., Hawkins, J., Whipple, C., Chuck, G., 2019. The regulatory landscape of a core maize domestication module controlling bud dormancy and growth repression. Nat. Commun. 10, 1–15. 10.1038/s41467-019-11774-w

Edgar, R., Domrachev, M., Lash, A.E., 2002. Gene Expression Omnibus: NCBI gene expression and hybridization array data repository. Nucleic Acids Res. 30, 207–210. 10.1093/nar/30.1.207

Emms, D.M., Kelly, S., 2019. OrthoFinder: phylogenetic orthology inference for comparative genomics. Genome Biol. 20, 238. 10.1186/s13059-019-1832-y

Engler, C., Youles, M., Gruetzner, R., Ehnert, T.M., Werner, S., Jones, J.D.G., Patron, N.J., Marillonnet, S., 2014. A Golden Gate modular cloning toolbox for plants. ACS Synth. Biol. 3, 839–843. 10.1021/sb4001504

Frei, D., Veekman, E., Grogg, D., Stoffel-Studer, I., Morishima, A., Shimizu-Inatsugi, R., Yates, S., Shimizu, K.K., Frey, J.E., Studer, B., Copetti, D., 2021. Ultralong Oxford Nanopore Reads Enable the Development of a Reference-Grade Perennial Ryegrass Genome Assembly. Genome Biol. Evol. 13, evab159. 10.1093/gbe/evab159

Gonzalez-Grandio, E., Pajoro, A., Franco-Zorrilla, J.M., Tarancon, C., Immink, R.G.H., Cubas, P., 2017. Abscisic acid signaling is controlled by a BRANCHED1/HD-ZIP i cascade in Arabidopsis axillary buds. Proc. Natl. Acad. Sci. U. S. A. 114, E245–E254. 10.1073/pnas.1613199114

Hajheidari, M., Huang, S.C., 2022. Elucidating the biology of transcription factor–DNA interaction for accurate identification of cis-regulatory elements. Curr. Opin. Plant Biol. 68, 102232. 10.1016/j.pbi.2022.102232

Hensel, G., Kastner, C., Oleszczuk, S., Riechen, J., Kumlehn, J., 2009. *Agrobacterium*-Mediated Gene Transfer to Cereal Crop Plants: Current Protocols for Barley, Wheat, Triticale, and Maize. Int. J. Plant Genomics 2009, e835608. 10.1155/2009/835608

Huang, D., Zheng, Q., Melchkart, T., Bekkaoui, Y., Konkin, D.J.F., Kagale, S., Martucci, M., You, F.M., Clarke, M., Adamski, N.M., Chinoy, C., Steed, A., McCartney, C.A., Cutler, A.J., Nicholson, P., Feurtado, J.A., 2020. Dominant inhibition of awn development by a putative zinc-finger transcriptional repressor expressed at the B1 locus in wheat. New Phytol. 225, 340–355. 10.1111/nph.16154

Hussien, A., Tavakol, E., Horner, D.S., Muñoz-Amatriaín, M., Muehlbauer, G.J., Rossini, L., 2014. Genetics of Tillering in Rice and Barley. Plant Genome 7, plantgenome2013.10.0032. 10.3835/PLANTGENOME2013.10.0032

Kerstens, M., Galinha, C., Hofhuis, H., Nodine, M., Pardal, R., Scheres, B., Willemsen, V., 2024. PLETHORA transcription factors promote early embryo development through induction of meristematic potential. Development 151, dev202527. 10.1242/dev.202527

Klasfeld, S., Roulé, T., Wagner, D., 2022. Greenscreen: A simple method to remove artifactual signals and enrich for true peaks in genomic datasets including ChIP-seq data. Plant Cell 34, 4795–4815. 10.1093/PLCELL/KOAC282

Kong, X., Wang, F., Geng, S., Guan, J., Tao, S., Jia, M., Sun, G., Wang, Z., Wang, K., Ye, X., Ma, J., Liu, D., Wei, Y., Zheng, Y., Fu, X., Mao, L., Lan, X., Li, A., 2022. The wheat AGL6-like MADS-box gene is a master regulator for floral organ identity and a target for spikelet meristem development manipulation. Plant Biotechnol. J. 20, 75–88. 10.1111/pbi.13696

Kosugi, S., Ohashi, Y., 2002. DNA binding and dimerization specificity and potential targets for the TCP protein family. Plant J. 30, 337–348. 10.1046/j.1365-313X.2002.01294.x

Kuijer, H.N.J., Shirley, N.J., Khor, S.F., Shi, J., Schwerdt, J., Zhang, D., Li, G., Burton, R.A., 2021. Transcript Profiling of MIKCc MADS-Box Genes Reveals Conserved and Novel Roles in Barley Inflorescence Development. Front. Plant Sci. 12, 1834. 10.3389/fpls.2021.705286

Lamesch, P., Berardini, T.Z., Li, D., Swarbreck, D., Wilks, C., Sasidharan, R., Muller, R., Dreher, K., Alexander, D.L., Garcia-Hernandez, M., Karthikeyan, A.S., Lee, C.H., Nelson, W.D., Ploetz, L., Singh, S., Wensel, A., Huala, E., 2012. The Arabidopsis Information Resource (TAIR): improved gene annotation and new tools. Nucleic Acids Res. 40, D1202–D1210. 10.1093/nar/gkr1090

Li, G., Kuijer, H.N.J., Yang, X., Liu, H., Shen, C., Shi, J., Betts, N., Tucker, M.R., Liang, W., Waugh, R., Burton, R.A., Zhang, D., 2021. MADS1 maintains barley spike morphology at high ambient temperatures. Nat. Plants 7, 1093–1107. 10.1038/s41477-021-00957-3

Li, H., 2013. Aligning sequence reads, clone sequences and assembly contigs with BWA-MEM. 10.48550/arXiv.1303.3997

Li, H., Durbin, R., 2010. Fast and accurate long-read alignment with Burrows–Wheeler transform. Bioinformatics 26, 589–595. 10.1093/BIOINFORMATICS/BTP698

Liller, C.B., Neuhaus, R., Von Korff, M., Koornneef, M., Van Esse, W., 2015. Mutations in barley row type genes have pleiotropic effects on shoot branching. PLoS ONE 10. 10.1371/journal.pone.0140246

Liu, H., Li, G., Yang, X., Kuijer, H.N.J., Liang, W., Zhang, D., 2020. Transcriptome profiling reveals phase-specific gene expression in the developing barley inflorescence. Crop J. 8, 71–86. 10.1016/j.cj.2019.04.005

Love, M.I., Huber, W., Anders, S., 2014. Moderated estimation of fold change and dispersion for RNA-seq data with DESeq2. Genome Biol. 15, 550. 10.1186/s13059-014-0550-8

Lu, Z., Marand, A.P., Ricci, W.A., Ethridge, C.L., Zhang, X., Schmitz, R.J., 2019. The prevalence, evolution and chromatin signatures of plant regulatory elements. Nat. Plants 5, 1250–1259. 10.1038/s41477-019-0548-z

Maher, K.A., Bajic, M., Kajala, K., Reynoso, M., Pauluzzi, G., West, D.A., Zumstein, K., Woodhouse, M., Bubb, K., Dorrity, M.W., Queitsch, C., Bailey-Serres, J., Sinha, N., Brady, S.M., Deal, R.B., 2018. Profiling of accessible chromatin regions across multiple plant species and cell types reveals common gene regulatory principles and new control modules. Plant Cell 30, 15–36. 10.1105/tpc.17.00581

Martín-Trillo, M., Grandío, E.G., Serra, F., Marcel, F., Rodríguez-Buey, M.L., Schmitz, G., Theres, K., Bendahmane, A., Dopazo, H., Cubas, P., 2011. Role of tomato BRANCHED1-like genes in the control of shoot branching. Plant J. 67, 701–714. 10.1111/j.1365-313X.2011.04629.x

Mascher, M., Wicker, T., Jenkins, J., Plott, C., Lux, T., Koh, C.S., Ens, J., Gundlach, H., Boston, L.B., Tulpová, Z., Holden, S., Hernández-Pinzón, I., Scholz, U., Mayer, K.F.X., Spannagl, M., Pozniak, C.J., Sharpe, A.G., Simková, H., Moscou, M.J., Grimwood, J., Schmutz, J., Stein, N., 2021. Long-read sequence assembly: a technical evaluation in barley. Plant Cell 33, 1888–1906. 10.1093/PLCELL/KOAB077

Niwa, M., Daimon, Y., Kurotani, K.I., Higo, A., Pruneda-Paz, J.L., Breton, G., Mitsuda, N., Kay, S.A., Ohme-Takagi, M., Endo, M., Araki, T., 2013. BRANCHED1 interacts with FLOWERING LOCUS T to repress the floral transition of the axillary meristems in Arabidopsis. Plant Cell 25, 1228–1242. 10.1105/tpc.112.109090

O’Malley, R.C., Huang, S.S.C., Song, L., Lewsey, M.G., Bartlett, A., Nery, J.R., Galli, M., Gallavotti, A., Ecker, J.R., 2016. Cistrome and Epicistrome Features Shape the Regulatory DNA Landscape. Cell 165, 1280–1292. 10.1016/j.cell.2016.04.038

Ouyang, S., Zhu, W., Hamilton, J., Lin, H., Campbell, M., Childs, K., Thibaud-Nissen, F., Malek, R.L., Lee, Y., Zheng, L., Orvis, J., Haas, B., Wortman, J., Buell, C.R., 2007. The TIGR Rice Genome Annotation Resource: improvements and new features. Nucleic Acids Res. 35, D883–D887. 10.1093/nar/gkl976

Patro, R., Duggal, G., Love, M.I., Irizarry, R.A., Kingsford, C., 2017. Salmon provides fast and bias-aware quantification of transcript expression. Nat. Methods 14, 417–419. 10.1038/nmeth.4197

Quinlan, A.R., Hall, I.M., 2010. BEDTools: a flexible suite of utilities for comparing genomic features. Bioinformatics 26, 841–842. 10.1093/bioinformatics/btq033

Ramsay, L., Comadran, J., Druka, A., Marshall, D.F., Thomas, W.T.B., MacAulay, M., MacKenzie, K., Simpson, C., Fuller, J., Bonar, N., Hayes, P.M., Lundqvist, U., Franckowiak, J.D., Close, T.J., Muehlbauer, G.J., Waugh, R., 2011. INTERMEDIUM-C, a modifier of lateral spikelet fertility in barley, is an ortholog of the maize domestication gene TEOSINTE BRANCHED 1. Nat. Genet. 43, 169–172. 10.1038/ng.745

Riaz, A., Alqudah, A.M., Kanwal, F., Pillen, K., Ye, L., Dai, F., Zhang, G., 2023. Advances in studies on the physiological and molecular regulation of barley tillering. J. Integr. Agric. 22, 1–13. 10.1016/j.jia.2022.08.011

Robinson, J.T., Thorvaldsdóttir, H., Winckler, W., Guttman, M., Lander, E.S., Getz, G., Mesirov, J.P., 2011. Integrative genomics viewer. Nat. Biotechnol. 29, 24–26. 10.1038/nbt.1754

Shaaf, S., Bretani, G., Biswas, A., Fontana, I.M., Rossini, L., 2019. Genetics of barley tiller and leaf development. J. Integr. Plant Biol. 61, 226–256. 10.1111/jipb.12757

Shang, Q., Wang, Y., Tang, H., Sui, N., Zhang, X., Wang, F., 2021. Genetic, hormonal, and environmental control of tillering in wheat. Crop J. 9, 986–991. 10.1016/j.cj.2021.03.002

Studer, A., Zhao, Q., Ross-Ibarra, J., Doebley, J., 2011. Identification of a functional transposon insertion in the maize domestication gene tb1. Nat. Genet. 43, 1160–1163. 10.1038/ng.942

Thiel, J., Koppolu, R., Trautewig, C., Hertig, C., Kale, S.M., Erbe, S., Mascher, M., Himmelbach, A., Rutten, T., Esteban, E., Pasha, A., Kumlehn, J., Provart, N.J., Vanderauwera, S., Frohberg, C., Schnurbusch, T., 2021. Transcriptional landscapes of floral meristems in barley. Sci. Adv. 7, 832–860. 10.1126/sciadv.abf0832

Törönen, P., Medlar, A., Holm, L., 2018. PANNZER2: a rapid functional annotation web server. Nucleic Acids Res. 46, W84–W88. 10.1093/nar/gky350

van Es, S.W., Muñoz-Gasca, A., Romero-Campero, F.J., González-Grandío, E., de los Reyes, P., Tarancón, C., van Dijk, A.D.J., van Esse, W., Pascual-García, A., Angenent, G.C., Immink, R.G.H., Cubas, P., 2023. A gene regulatory network critical for axillary bud dormancy directly controlled by Arabidopsis BRANCHED1. New Phytol. n/a. 10.1111/nph.19420

van Esse, G.W., Walla, A., Finke, A., Koornneef, M., Pecinka, A., von Korff, M., 2017. Six-Rowed Spike3 (VRS3) Is a Histone Demethylase That Controls Lateral Spikelet Development in Barley. Plant Physiol. 174, 2397–2408. 10.1104/pp.17.00108

Vogel, J.P., Garvin, D.F., Mockler, T.C., Schmutz, J., Rokhsar, D., Bevan, M.W., Barry, K., Lucas, S., Harmon-Smith, M., Lail, K., Tice, H., Schmutz (Leader), J., Grimwood, J., McKenzie, N., Bevan, M.W., Huo, N., Gu, Y.Q., Lazo, G.R., Anderson, O.D., Vogel (Leader), J.P., You, F.M., Luo, M.-C., Dvorak, J., Wright, J., Febrer, M., Bevan, M.W., Idziak, D., Hasterok, R., Garvin, D.F., Lindquist, E., Wang, M., Fox, S.E., Priest, H.D., Filichkin, S.A., Givan, S.A., Bryant, D.W., Chang, J.H., Mockler (Leader), T.C., Wu, H., Wu, W., Hsia, A.-P., Schnable, P.S., Kalyanaraman, A., Barbazuk, B., Michael, T.P., Hazen, S.P., Bragg, J.N., Laudencia-Chingcuanco, D., Vogel, J.P., Garvin, D.F., Weng, Y., McKenzie, N., Bevan, M.W., Haberer, G., Spannagl, M., Mayer (Leader), K., Rattei, T., Mitros, T., Rokhsar, D., Lee, S.-J., Rose, J.K.C., Mueller, L.A., York, T.L., Wicker (Leader), T., Buchmann, J.P., Tanskanen, J., Schulman (Leader), A.H., Gundlach, H., Wright, J., Bevan, M., Costa de Oliveira, A., da C. Maia, L., Belknap, W., Gu, Y.Q., Jiang, N., Lai, J., Zhu, L., Ma, J., Sun, C., Pritham, E., Salse (Leader), J., Murat, F., Abrouk, M., Haberer, G., Spannagl, M., Mayer, K., Bruggmann, R., Messing, J., You, F.M., Luo, M.-C., Dvorak, J., Fahlgren, N., Fox, S.E., Sullivan, C.M., Mockler, T.C., Carrington, J.C., Chapman, E.J., May, G.D., Zhai, J., Ganssmann, M., Guna Ranjan Gurazada, S., German, M., Meyers, B.C., Green (Leader), P.J., Bragg, J.N., Tyler, L., Wu, J., Gu, Y.Q., Lazo, G.R., Laudencia-Chingcuanco, D., Thomson, J., Vogel (Leader), J.P., Hazen, S.P., Chen, S., Scheller, H.V., Harholt, J., Ulvskov, P., Fox, S.E., Filichkin, S.A., Fahlgren, N., Kimbrel, J.A., Chang, J.H., Sullivan, C.M., Chapman, E.J., Carrington, J.C., Mockler, T.C., Bartley, L.E., Cao, P., Jung, K.-H., Sharma, M.K., Vega-Sanchez, M., Ronald, P., Dardick, C.D., De Bodt, S., Verelst, W., Inzé, D., Heese, M., Schnittger, A., Yang, X., Kalluri, U.C., Tuskan, G.A., Hua, Z., Vierstra, R.D., Garvin, D.F., Cui, Y., Ouyang, S., Sun, Q., Liu, Z., Yilmaz, A., Grotewold, E., Sibout, R., Hematy, K., Mouille, G., Höfte, H., Michael, T., Pelloux, J., O’Connor, D., Schnable, J., Rowe, S., Harmon, F., Cass, C.L., Sedbrook, J.C., Byrne, M.E., Walsh, S., Higgins, J., Bevan, M., Li, P., Brutnell, T., Unver, T., Budak, H., Belcram, H., Charles, M., Chalhoub, B., Baxter, I., The International Brachypodium Initiative, Principal investigators, DNA sequencing and assembly, Pseudomolecule assembly and BAC end sequencing, Transcriptome sequencing and analysis, Gene analysis and annotation, Repeats analysis, Comparative genomics, Small RNA analysis, Manual annotation and gene family analysis, 2010. Genome sequencing and analysis of the model grass Brachypodium distachyon. Nature 463, 763–768. 10.1038/nature08747

Waddington, S.R., Cartwright, P.M., Wall, P.C., 1983. A Quantitative Scale of Spike Initial and Pistil Development in Barley and Wheat. Ann. Bot. 51, 119–130. 10.1093/oxfordjournals.aob.a086434

Wang, H., Chen, W., Eggert, K., Charnikhova, T., Bouwmeester, H., Schweizer, P., Hajirezaei, M.R., Seiler, C., Sreenivasulu, N., von Wirén, N., Kuhlmann, M., 2018. Abscisic acid influences tillering by modulation of strigolactones in barley. J. Exp. Bot. 69, 3883–3898. 10.1093/jxb/ery200

Wang, H., Seiler, C., Sreenivasulu, N., Von Wirén, N., Kuhlmann, M., 2022. INTERMEDIUM-C mediates the shade-induced bud growth arrest in barley. J. Exp. Bot. 73, 1963–1977. 10.1093/jxb/erab542

Wang, R.-L., Stec, A., Hey, J., Lukens, L., Doebley, J., 1999. The limits of selection during maize domestication. Nature 398, 236–239. 10.1038/18435

Yaffe, H., Buxdorf, K., Shapira, I., Ein-Gedi, S., Moyal-Ben Zvi, M., Fridman, E., Moshelion, M., Levy, M., 2012. LogSpin: A simple, economical and fast method for RNA isolation from infected or healthy plants and other eukaryotic tissues. BMC Res. Notes 5, 1–8. 10.1186/1756-0500-5-45/FIGURES/3

Zhang, Y., Liu, T., Meyer, C.A., Eeckhoute, J., Johnson, D.S., Bernstein, B.E., Nussbaum, C., Myers, R.M., Brown, M., Li, W., Shirley, X.S., 2008. Model-based analysis of ChIP-Seq (MACS). Genome Biol. 9, 1–9. 10.1186/gb-2008-9-9-r137

Zhang, Y., Shen, C., Li, G., Shi, Jin, Yuan, Y., Ye, L., Song, Q., Shi, Jianxin, Zhang, D., 2024. MADS1-regulated lemma and awn development benefits barley yield. Nat. Commun. 15, 301. 10.1038/s41467-023-44457-8

Zhang, Yuyun, Li, Z., Zhang, Yu’e, Lin, K., Peng, Y., Ye, L., Zhuang, Y., Wang, M., Xie, Y., Guo, J., Teng, W., Tong, Y., Zhang, W., Xue, Y., Lang, Z., Zhang, Yijing, 2021. Evolutionary rewiring of the wheat transcriptional regulatory network by lineage-specific transposable elements. Genome Res. 31, 2276–2289. 10.1101/gr.275658.121

Zhang, Z., Zhao, P., Wang, X., Wang, H., Zhai, Z., Zhao, X., Xing, L., Qi, Z., Shang, Y., 2024. Identification and map-based cloning of long glume mutant gene lgm1 in barley. Mol. Breed. 44, 3. 10.1007/s11032-024-01448-x

Zhou, W., Shi, H., Wang, Z., Huang, Y., Ni, L., Chen, X., Liu, Yan, Li, H., Li, C., Liu, Yaxi, 2024. Identification of highly repetitive barley enhancers with long-range regulation potential via STARR-seq. Genomics Proteomics Bioinformatics qzae012. 10.1093/gpbjnl/qzae012

Zwirek, M., Waugh, R., McKim, S.M., 2019. Interaction between row-type genes in barley controls meristem determinacy and reveals novel routes to improved grain. New Phytol. 221, 1950–1965. 10.1111/nph.15548

